# Supplemental Dietary Nano Zinc Can Impart Higher Antioxidant Status by Modulating Cu-Zn SOD Expression

**DOI:** 10.1101/2024.09.21.614296

**Authors:** Amrita Tah, Aruna Pal, Debasis De, P.N. Chatterjee

## Abstract

Zinc seems to be the most critical micronutrients dietary adequacy of which ensures optimum health and body defence. Considering its immunomodulation potency, zinc is often used for dietary fortification more particularly in face of challenges. However indiscriminate use of zinc as therapeutic agent often leads to secondary deficiencies of other critical nutrients due to the unwanted antagonism with other interacting nutrients available in gut. In the present study we targeted to devise a nano-structured zinc which will remain inert to other dietary micronutrients present in gut to ensure its high bioavailability. An environment benign colloidal chemistry route was employed for the synthesis of nano-dimensional zinc oxide. The microanalytical characterizations revealed that the apparent particle size of our nano-Zinc oxide ranged between 30-40 nm. The as-synthesized ZnO-NP was used for dietary fortification and the MTT assay confirmed the safe limit for its dietary inclusion. The antioxidant potential of the synthesized nano-zinc was evaluated in milk fish (*Chanos chanos*) considering it as a model organism. We have employed Cu-Zn SOD gene as molecular marker for antioxidant assessment with integral zinc binding sites. We have characterized Cu-Zn SOD gene in *Chanos chanos* for the first time and identified certain important zinc binding sites present in the Cu-Zn SOD. A significantly better expression profile of Cu-Zn SOD was observed among the fish fed dietary ZnO-NP and the best effect was observed when the fish feed was fortified with 20 ppm ZnO-NP. The outcome of this study ensures the higher bioavailability of the synthesized ZnO-NP to be assimilated into Cu-Zn SOD, which in turn imparts higher body antioxidant. Nano zinc being inert, may directly bind through the zinc binding sites of Cu-Zn SOD molecules thereby leads to its better expression and more antioxidant status through molecular interaction with other molecules through *longevity regulating pathway* as explored by the String and KEGG pathway analysis carried out in the present investigation.

## Introduction

The recent advances in nanobiotechnology have reshaped the scientific innovations with newer molecular insights to understand the basis of different biological processes. The higher surface to charge ratio offers the nanoscale materials some unique properties to carry out the desired function. Researches are in vogue to design different varieties of nanostructures for precising the production of vaccines, drugs, feed additives, nutrient delivery, disease treatmentetc (1).

Micronutrients play a crucial role in growth, production and reproduction and body defence of all categories of animals including fish and it needs a regular intake to meet the physiological deficiencies. Mere fulfilling the requirement of a mineral might not assure its adequacy at cellular level. The bioavailability of minerals is the most challenging issue (2) in the science of nutrition. Mineral’s bioavailability is mostly regulated by its source and the presence of other interacting micro-nutrients present in gut. In recent days different varieties of nano-minerals have been synthesized which appeared to be highly bioavailable and hence has least interaction with other nutrients available in gut. Supplemental nano-minerals might have the ability to enhance the growth and immunity (3). The synthesized nanoparticle is might be more efficient than its bulk counterparts. Several metal oxide nanoparticles have proved their antimicrobial activities against a wide range of bacteria and fungi (4).

Zinc is the second most abundant trace element of the body but it cannot be stored in the bodythat is why regular dietary intake of zinc is needed to avoid any subclinical deficiency (5, 6). Zinc plays a vital role in prostaglandin metabolism and provides structural rigidity to the nucleoproteins (7). Zinc act as an essential part of about 20 metalloenzymes that includes alkaline phosphatase, alcohol dehydrogenase, carbonic anhydrase etc. Besides this zinc plays critical role in growth performance, fertility, immune function, wound healing, oxidative stress maintenance etc (5, 8, 9). Zinc oxide and nano-zinc oxide have the same chemical structure which suggests same zinc and oxygen ratio, but the nano-sized atoms have wider energy level confinement and due to size variation, it becomes less interactive with other bulk size molecules available in the gut (10). The higher bioavailability of nano-structures ensures its high accumulation at tissue level and hence ZnO-NP appeared to be more efficient for imparting high antioxidant status in fish (11). The accumulation of zinc in the tissue in sufficient amount ensures highercatalase, SOD and glutathione peroxidase activity (12). The increased amount of SOD and GPx enhance the activity of NADPH oxidase which is accountable for hunting of the superoxide anions (13). Zinc serves as a co-factor and active component for the SOD (14).

Fish employs three different isoforms of SOD and they are characterized by their metal co-factors and cellular localization. Three isoforms of SOD include: Cu-Zn SOD which present in the cytosol (SOD1); Mn-SOD which is located in mitochondria (SOD2); and extracellular Cu-Zn SODthat is located in the extracellular matrix of tissue (SOD3) (15, 16, 17). The SOD plays crucial role in development andalso take part in the maintenance of homeostatic balance between innate immune response and antioxidant defences. SOD1 is a homodimer and this homodimer is formed byconnecting two identical subunits through both hydrophobic interactions by the residues Val^6^, Val^8^,Ile^18, 114^IGR^116^, and ^149^VIGIAQ^116^and electrostatic interactions, by the residues Glu^130^, Glu^131^, Lys^137^, and Thr^138^(18).

In recent days animal nutritionists are trying to develop zinc-based feed additives with an aim to improve the antioxidant status of fish for ensuring higher body defense. Being more bioavailable and least antagonistic to other micronutrients, ZnO-NPmight serve as a feasible alternative. Fish are continuously being exposed to a plethora of stresses. The fishes at their early life are more vulnerable to environmental extremes. The embryonic and larval development require dynamic cellular activities with high oxygen/energy demands, involving changes in metabolism, physiology, and body architecture all in turn leads to increase in ROS production. This is quite critical and often leads to damage of hepatocytes andthereby increased the mortality rate (19, 20). Some sporadic experiments on dietary zinc for imparting higher antioxidant status have been carried out in freshwater fish but scanty reports are available on brackish water fish. The culture of Milk Fish (*Chanos chanos)* is ever increasing in coastal belt of India and in some parts of southeast Asia and Taiwan (21) considering its delicacy coupled with its elite nutrient profile. It is an important euryhaline finfish that can be cultured in fresh water, brackishwater or seawater (21, 22).

The present experiment was aimed to compare the effect of formulated diets fortified with different sources of zinc (Inorganic, organic and nano-zinc) to evaluate its role on Cu-Zn SOD gene expression which is essential for imparting higher antioxidant status in *Chanos chanos* (milk fish).

## Materials and Methods

The present study was aimed at exploring the antioxidant mechanism offered by feeding of ZnO-NP fortified diet in milk fish (*Chanoschanos*) model. The nano-zinc have been synthesized and used as a dietary supplement. We have employed Cu-Zn SOD as the molecular marker for the detection of the antioxidant status. We have characterized Cu-Zn SOD gene in *Chanoschanos* for the first time and assessed the differential mRNA expression level in liver with respect to inorganic zinc, organic zinc, 10 ppm ZnO-NP, 20 and 40 ppm ZnO-NP.

### Chemical synthesis of dietary nano-zinc

The nano-dimensional zinc oxide particles (ZnO-NPs) have been synthesized in our laboratory by adopting the colloidal chemistry route as stated earlier (23). An aqueous solution of thecationic surfactant, cetyltrimethyl ammonium bromide (CTAB) (12 mM) was added (10 mL) dropwise to zinc acetate (6 mM) aqueoussolution (25 mL) with continuous stirring for 30 min. Subsequently, 4 mL ammonia (30%) was added dropwise to this mixture, and the solution was stirred for 15 min. The whole content was then transferred into stoppered glass tubes (10 mL capacity) and kept in oven at 100 °C for 24 h followed by its cooling and centrifugation. The precipitate obtained was firstwashed with water and then with ethanol repetitively by centrifugation. The final white precipitate was dried for 2 h at 60 °C in a hot air oven to obtain the soft powder of ZnO-NPs. The as synthesized nanostructures were characterized microanalytically and then preserved in sealed vials for further use.

### Characterization of ZnO Nanoparticles

The as synthesized ZnO-NPs werecharacterized first through field-emission scanning electron microscope (FESEM; SUPRA 55 VP 4132 Carl Zeiss, Germeny) at Indian Institute of Science Education and Research Kolkata, India. The elemental composition of the synthesized nanoparticles was performed by energy-dispersive X-ray (EDX) (SUPRA 55 VP 4132 Carl Zeiss, Oberkochen, Germany) at Indian Institute of Science Education and Research Kolkata, India. The powder X-ray diffraction (XRD) (Bruker D8 Advance, Karlsruhe, Germany) was performed at Central Instrumentation Facility, Kalyani University, India for identification of purity and quantitative analysis of ZnO-NPs within range from 30° to 80° Cu *K*α radiations (*k* =0.15406 nm).

The biosafety of the as synthesized nano-zinc was assessed through MTT assay (Chatterjee et al., 2023-conference paper State DST, Govt. of West Bengal) and included within the safe limit for dietary fortification.

### Experimental animals and dietary allocation

Disease-free healthy motile active *Chanoschanos* fries (0.35 ± 0.05 g) were procured from the pond of ICAR-Central Institute of Brakishwater Aquaculture (CIBA), Kakdwip Research Centre and were acclimatized for 15 days in 600 L fiber-reinforced plastic (FRP) tanks. They were fed twice daily (morning and evening) at 10% of their body weight. The waste was siphoned out and 80% waterwas exchanged at every three days intervals to ensure the proper living condition for the fish with least stress (24). No mortality was noticed during the acclimatization period. The fishes were observed for mortality daily throughout the experimental feeding trial and the dead fishes were removed immediately. The water quality parameters (pH, dissolved oxygen, ammonia-nitrogen etc) were monitored on a regular basis.

After initial acclimatization, a total of 540 fish were randomly allocated into six treatments (Table 1) each having three replicates and each replicate contains 30 fish fries. The fishes of T1 fed the basal diet without any zinc supplementation; the fishes belonged to T2 and T3 fed the basal diet supplemented with 20 ppm inorganic zinc oxide (ZnO) and organic-zinc (Zn-amino acid complex) (ZnA) respectively; and the fishes of the other three treatments (T4, T5, T6) were supplemented with 10, 20 and 40 ppm nano-zinc oxide (ZnO-NP) respectively as depicted in Table 1. The basal feed remained isonitrogenous and isocaloric throughout the feeding trial across the treatment groups.

**Table 1.** Experimental design.

| Group | Replicate | Dietary treatment |
| --- | --- | --- |
| T1 | 3 | Basal feed (Without zinc) |
| T2 | 3 | Basal feed + 20ppm inorganic zinc |
| T3 | 3 | Basal feed + 20ppm organic zinc |
| T4 | 3 | Basal feed + 10ppm Nano zinc |
| T5 | 3 | Basal feed + 20ppm Nano zinc |
| T6 | 3 | Basal feed + 40ppm Nano zinc |

The feeding trial was continued for 120 days and followed by a digestibility trial extending for 7 days. At the end of feeding trial, three fishes from each replicate were sacrificed by a registered veterinarian (PNC) and the liver was collected aseptically and stored in a sterile vial by keeping submerged in RNA-later solution.

### Characterization of Cu-Zn SOD Gene in *Chanoschanos*

The total RNA was isolated from the liver by Trizol method and subsequently converted to cDNA as per the standard protocol developed in our lab (25, 26, 27, 28).

### Materials

The Taq DNA polymerase, 10X buffer and dNTP were purchased from Invitrogen; the SYBR Green qPCR Master Mix (2X) was obtained from Thermo Fisher Scientific Inc. (PA, USA); the L-Glutamine (Glutamax 100x) was purchased from Invitrogen corp., (Carlsbad, CA, USA); the Penicillin-G and streptomycin were obtained from Amresco (Solon, OH, USA);the Filters (Millex GV. 0.22 µm) werepurchased from Millipore Pvt. Ltd., (Billerica, MA, USA). All other reagents were molecular biology grade and analytical.

### Synthesis, Confirmation of cDNA and PCR Amplification of gene

The reaction mixture of 20µlwas consists of 5μg of total RNA, 0.5μg of oligo dT primer (16– 18mer), 40U of Ribonuclease inhibitor, 10M of dNTP mix, 10mM of DTT, and 5U of MuMLV reverse transcriptase in reverse transcriptase buffer. There was gentle mixing of reaction mixture and incubated the reaction mixture at 37°C for 1 hour. The reaction was sat 70°C by heating it at 70º C for 10 minutes and keep it in chilledice. The integrity of the cDNA was checked by PCR. The primers have been listed in Table 2. 25μL reaction mixture contained 80–100ng cDNA, 3.0μL 10X PCR assay buffer, 0.5μL of 10mM dNTP, 1U Taq DNA polymerase, 60ng of each primer (as in Table 2), and 2mM MgCl_2_. PCR-reactions were carried out in a thermocycler (PTC-200, MJ Research, USA) with cycling conditions as, initial denaturation at 94°C for 3min, denaturation at 94°C for 30sec, annealing temperature 60º C for 35sec, and extension at 72°C for 3min was carried out for 35 cycles followed by final extension at 72°C for 10 min.

**Table 2:**
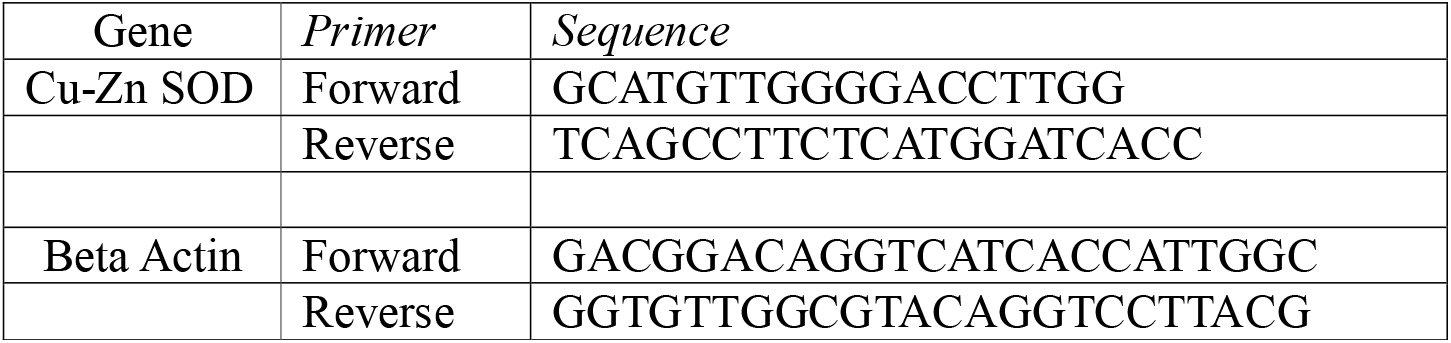
PCR primers for differential mRNA expression profiling. (35, 36);

The amplified products were sequenced with Sanger sequencing. Further *in silico analysis* were conducted with the Cu-Zn SOD gene sequences and submitted to public domain.

### Study of Predicted gene Using Bioinformatics Tools

The predicted peptide sequence of genes of milk fish was derived by Edit sequence (Lasergene Software, DNASTAR). Prediction of the signal peptide of the genes were conducted using the software (Signal P 3.0 Sewer-prediction results, Technical University of Denmark). Analysis of O-linked glycosylation sites was carried out using NetOGlyc 4 server (http://www.expassy.org/), whereas the N-linked glycosylation site was detected by NetNGlyc 1.0 software (http://www.expassy.org/). Regions for alpha-helix and beta-sheet were predicted using NetSurfP-Protein Surface Accessibility and Secondary Structure Predictions, Technical University of Denmark (29).

### Three-dimensional structure prediction and Model quality assessment

With highest sequence id the templates was possessed andentities with our target template were identified by using PSI-BLAST (http://blast.ncbi.nlm.nih.gov/Blast). The homology modeling was used to build a 3D structure based on homologous template structures using PHYRE2 server (30). The 3D structures were visualized by PyMOL(http://www.pymol.org/) which is an open-source molecular visualizationtool. Subsequently, the three dimensional model was generated using PyMoL tool. For controlling energy minimization the Swiss PDB Viewer was employed. The stereochemical quality assessment and structural evaluation of predicted model was carried out by using the SAVES (Structural Analysis and Verification Server), which is an integrated server (http://nihserver.mbi.ucla.edu/SAVES/). The ProSA (Protein Structure Analysis) webserver (https://prosa.services.came.sbg.ac.at/prosa) was used for refinement and validation of protein structure (31). The ProSA was used for checking model structural quality with potential errors and the program shows a plot of its residue energies and Zscores which determine the overall quality of the model. The solvent accessibility surface area of the genes was generated by using NetSurfPserver(http://www.cbs.dtu.dk/services/NetSurfP/)(29). It analyse relative surface accessibility of protein, Z-fit score, Alpha-Helix probability, probability for beta-strand and coil score, etc. TM align software was used for the alignment of 3 D structure of IR protein for different species and RMSD estimation to assess the structural differentiation (32). The Provean analysis was conducted to assess the deleterious nature of the mutant amino acid. PDB structure for 3D structural prediction of gene for milk fish was carried out through PHYRE software38. Protein-protein interaction have been studied through String analysis(33).

### Differential mRNA expression profiling of Cu-Zn SOD gene in *Chanoschanos*

#### Real time PCR

Total RNA was estimated from liver of milk fish from six treatments group by Trizol method. First strand cDNA was synthesised by the process of reverse transcriptase polymerase chain reaction (rt-PCR) in the automated temperature-maintained thermocycler machine. As reverse transcriptase enzymeM-MLVRT (200 u/µl) was used. The primer was obtained from a published journal (25). The primers used are listed in Table2. RNA of equal amount (quantified by Qubit fluorometer, Invitrogen), wherever applicable, were used for cDNA preparation (Superscript III cDNA synthesis kit; Invitrogen). All qRT-PCR reactions were conducted on ABI 7500 fast system. Each reaction consisted of 2 µl cDNA template, 5 µl of 2X SYBR Green PCR Master Mix, 0.25 µl each of forward and reverse primers (10 pmol/µl) and nuclease free water to make up final volume of 10 µl. Each sample was run in triplicate. Analysis of real-time PCR (qRT-PCR) was performed by delta-delta-Ct (ΔΔCt) method (25, 26, 28, 34).

The entire reactions were performed in triplicate (as per MIQE Guidelines) and experiment repeated twice, in 20µl reaction volume, using FastStart Essential DNA Green Master (Himedia) on ABI 7500 system.

### Statistical analysis

The descriptive statistical analysis was carried out through SYSTAT package for the expression level analysed through real time PCR with mean and standard error. It was presented accordingly in graph. The estimation of Expression level with real time PCR was carried out as 2^-ΔΔCt^.

## Result

### Microanalytical characterization of the synthesized nano-zinc

The Field-emission scanning electron microscopy (FESEM) study of the nano dimensional zinc oxide synthesizedin the present study revealed the bundled rod-like nanostructure. Each bundle is made up with concentrically grown spindle like nanostructures. The spindle like nanostructure seems to act as the growth unit of the bundled aggregates.The cationic surfactant CTAB might have controlled the shape and determined the one-dimensional structure of the synthesized nano-Zinc oxide.

Followed by FESEM, the synthesized nano-dimensional zinc oxide particles were subjected to Energy-dispersive X-ray (EDX) analysis which confirmed the presence of both zinc and oxygen molecule in the synthesized zinc oxide nano-structures. A powder XRD (PXRD) study was also carried out to assess the purity of the synthesized ZnO-NPs. ThePXRD pattern of the as-synthesized ZnO nanoparticles indicated thatthe synthesized nano particle is puredevoid of any contamination.

### Characterization of Cu-Zn SOD gene in *Chanoschanos*

#### Characterization of the Cu-Zn SOD genes: In silico studies and identification of important domains

Cu-Zn SOD gene has been employed in the current study as an important molecular marker for assessing the oxidative stress. Cu-Zn SOD gene has been characterized and predicted 3D structure for the peptide sequence is visualized (Fig 2).The 3D structure revealed that Cu-Zn SOD of *chanoschanos*is consists of beta barrel.The four zinc binding sites has been detected at amino acid position 64 (Fig 2, Green sphere), amino acid position 72 (Fig 2, Red sphere), amino acid position 81 (Fig 2, Chocolate sphere) and amino acid position 84 (Fig 2, Blue sphere). Secondary structure of Cu-Zn SOD gene of *Chanoschanos* is depicted in Fig 3.

**Figure 1.**
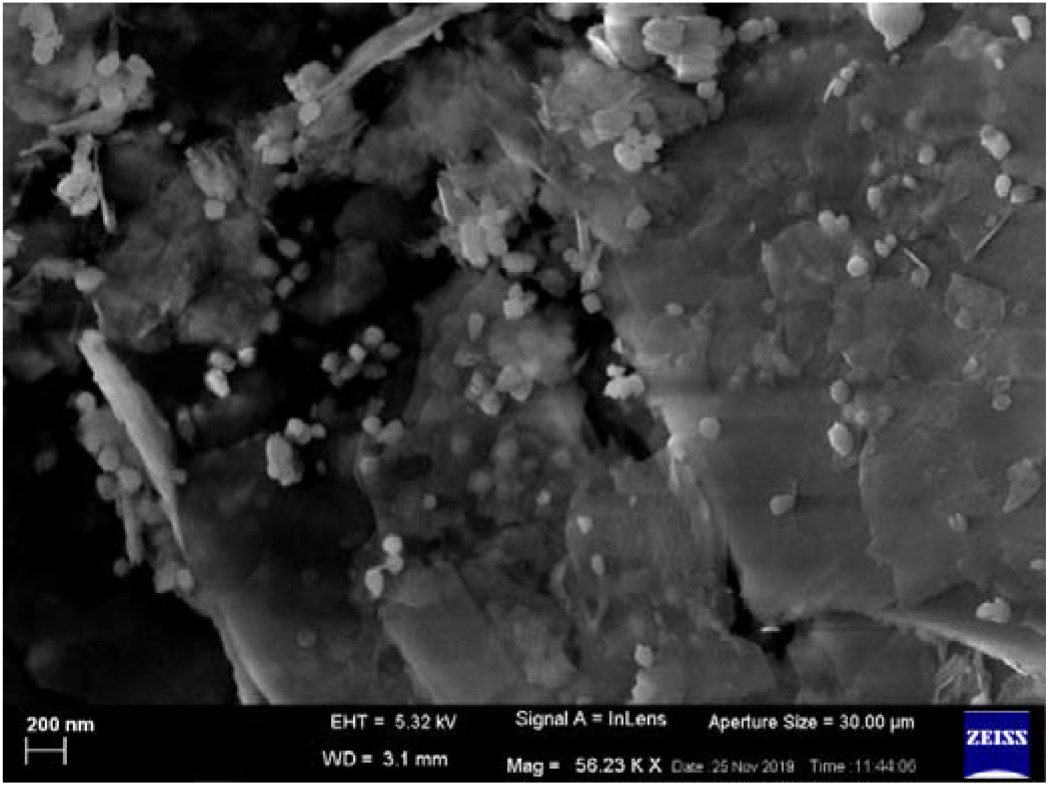
Field-emission scanning electron microscopy image of nano-Zinc oxide.

**Fig 2.**
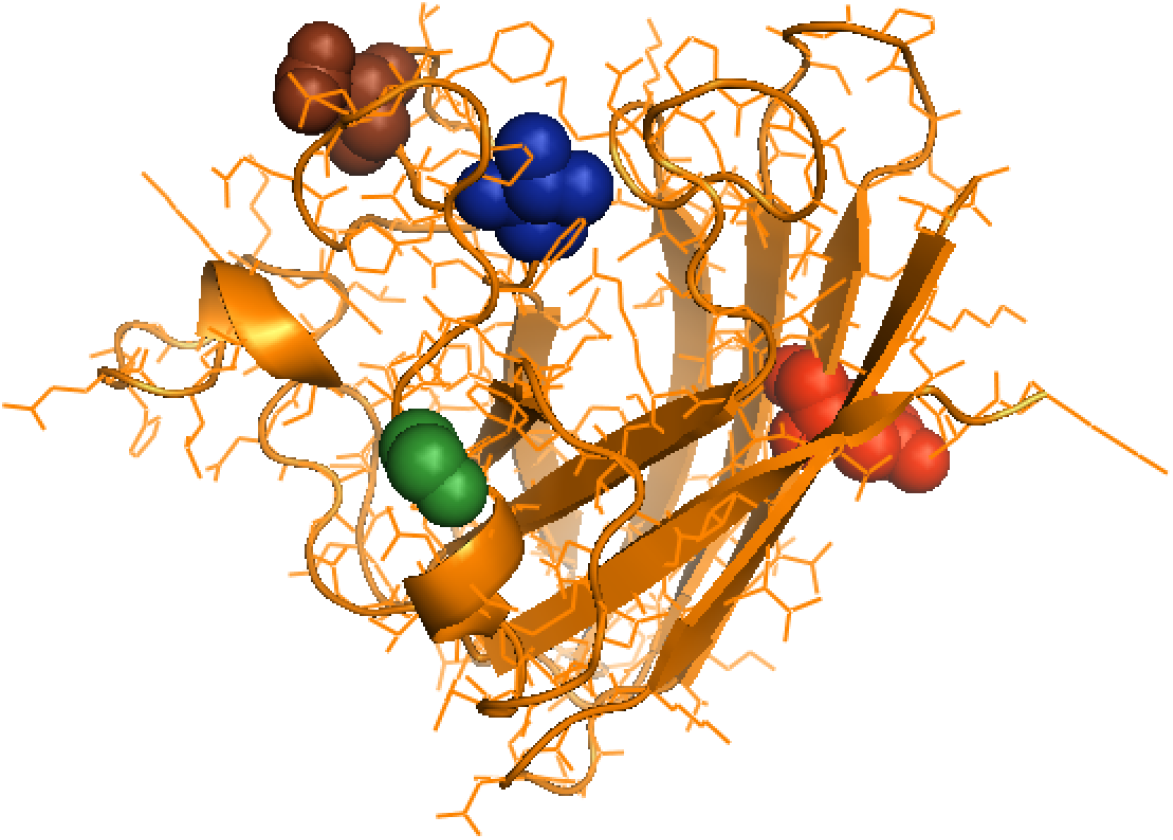
3D structure for Cu-Zn SOD in *Chanoschanos* with the domains for zinc binding sites (Zinc binding site aa 64: Green sphere; Zinc binding site aa 72: Red sphere; Zinc binding site aa 81: Chocolate sphere; Zinc binding site aa 84: Blue sphere)

**Fig 3.**
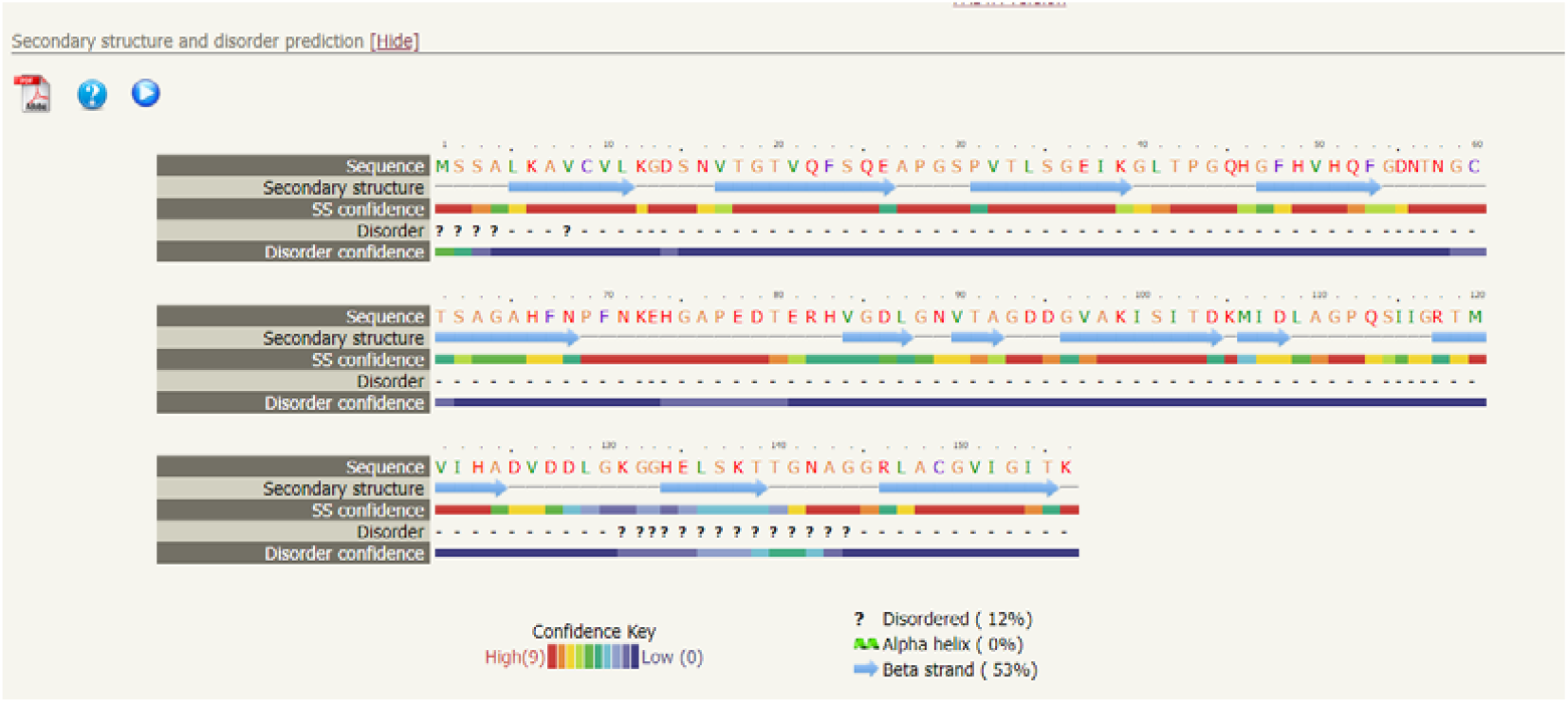
Secondary structure for Cu-Zn SOD gene for *Chanoschanos*.

### String Analysis

The string analysis revealed that Cu-Zn SOD interacts with many other genes in a complex biological array. These genes include TRX1, TRX2, TSA1, CTT1, GLR1, CTA1, SOD2, CCS1, COX17, ATX1 as per their interaction score (Fig 4 and Fig 5).

**Fig 4.**
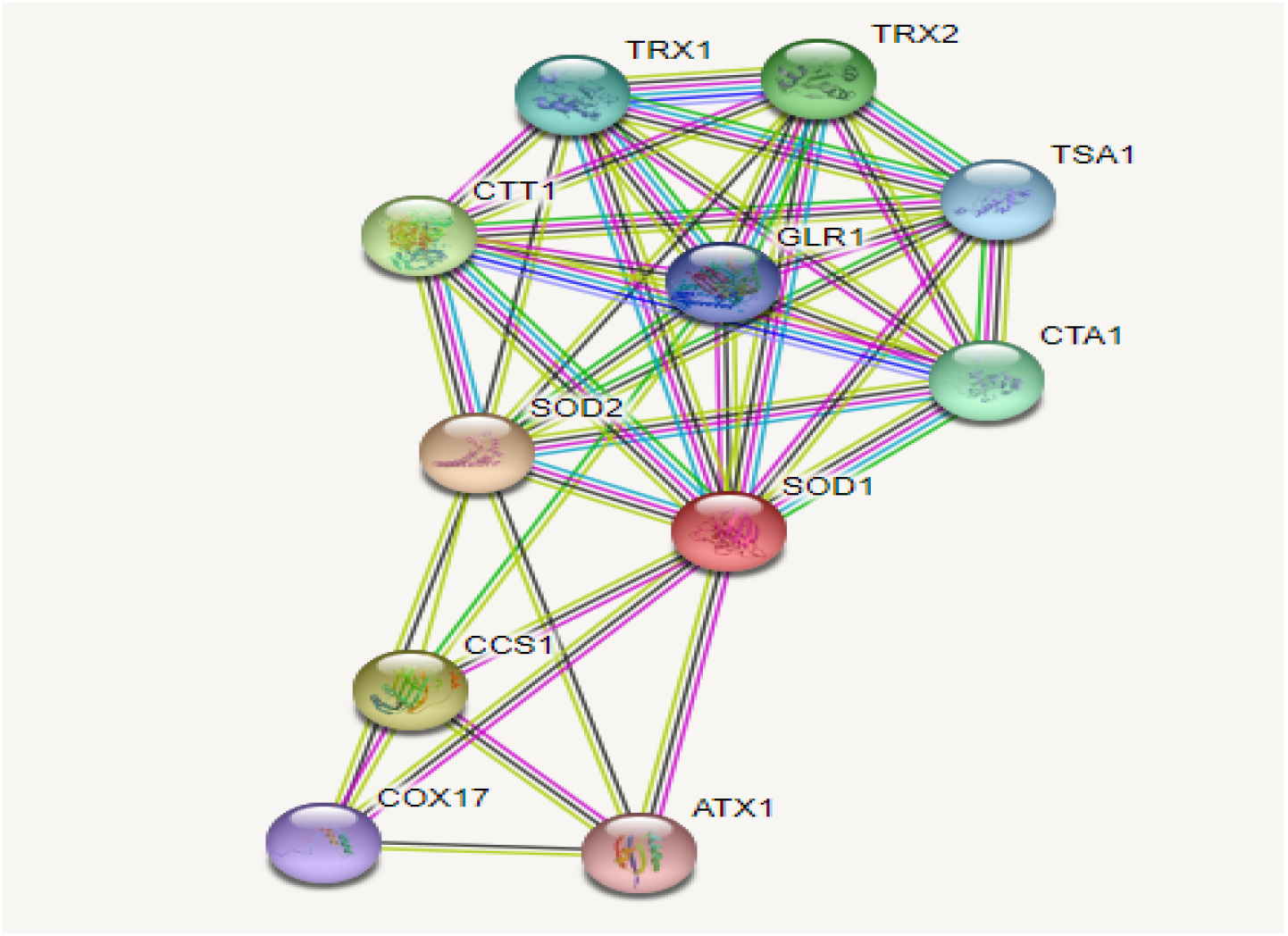
String analysis for Cu-Zn SOD gene for *Chanos chanos*.

**Fig 5.**
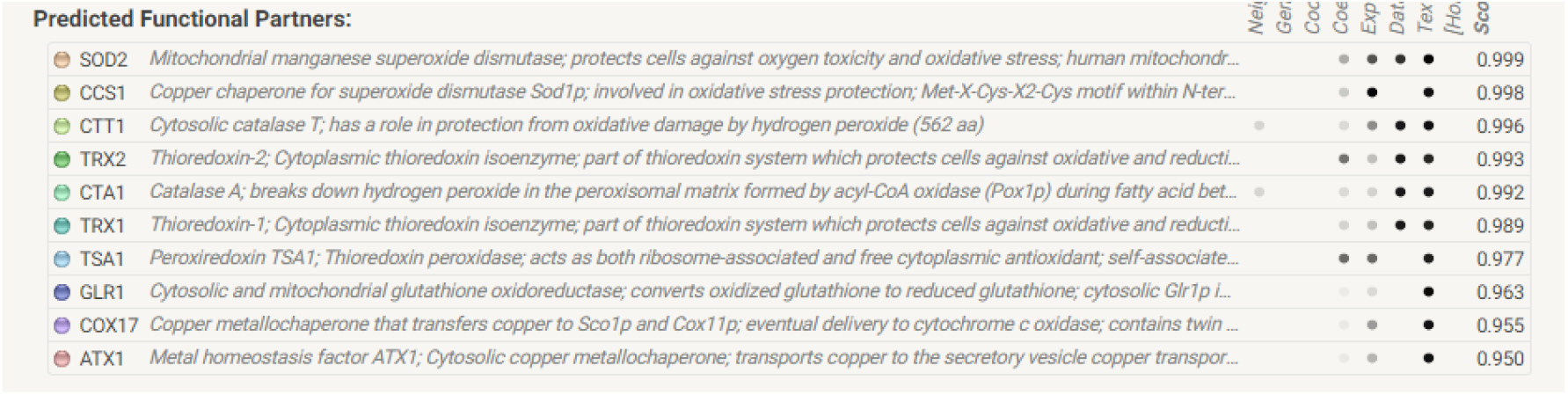
Interacting genes for *Chanos chanos* with Cu-Zn SOD

### KEGG Analysis

The KEGG analysis explored the role of Cu-Zn SOD in longevity regulating pathway which has been described in Fig 6. This outcome of the KEGG established the antioxidant effect of Cu-Zn SOD through diverse pathway.

**Fig 6.**
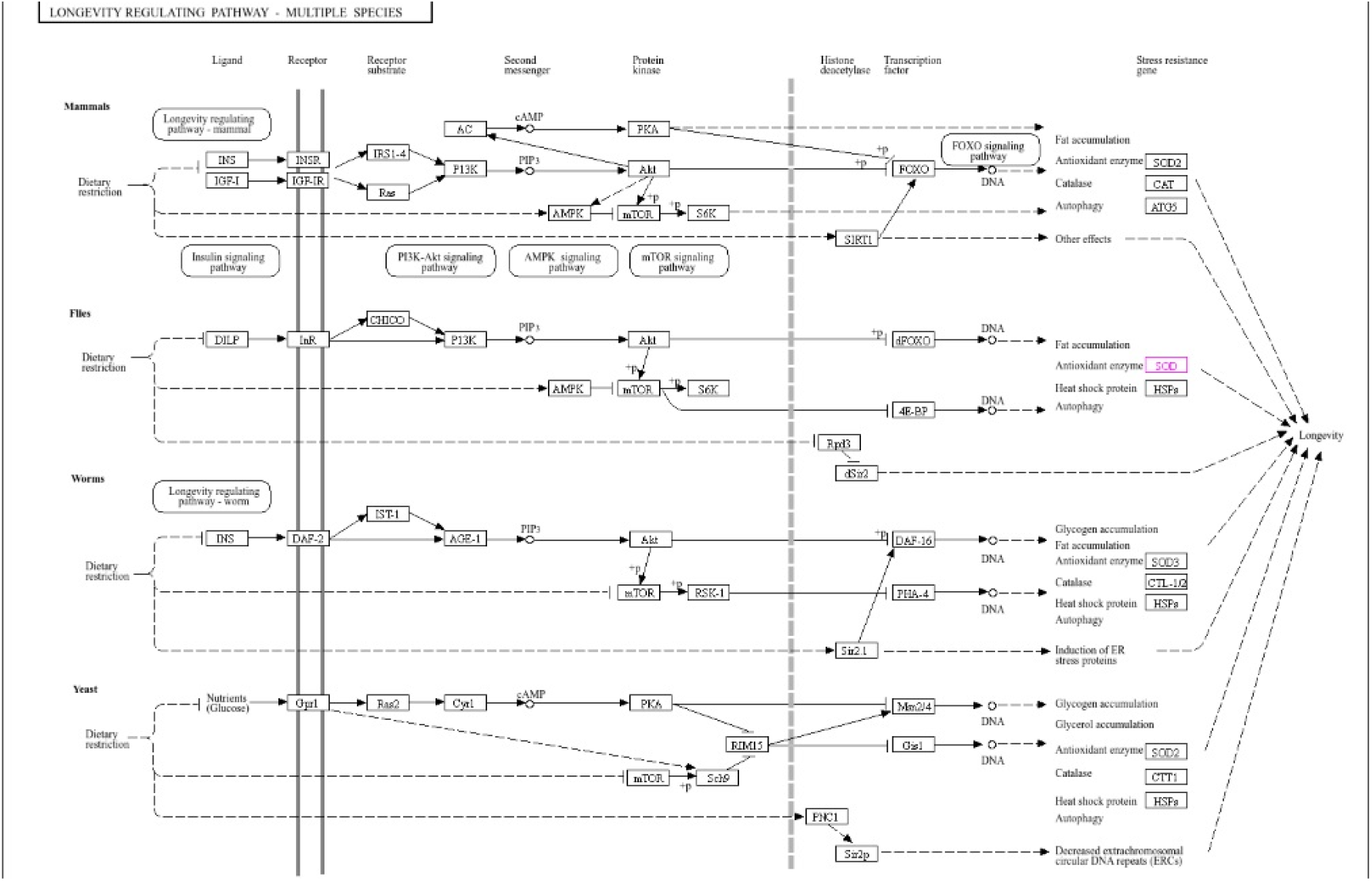
KEGG analysis of Cu-Zn SOD that shows antioxidant pathway through longevity regulating pathway.

### Differential mRNA expression profiling of the gene

In the present investigation the Cu-Zn SOD gene was observed to be upregulated significantly in treatment T5 (20 ppm ZnO-NP supplemented group) compared to other treatments (Fig7). Lowest expression was observed among the organic zinc supplemented fish. The nano-zinc fed fish depicted a higher Cu-Zn SOD expression even when fed @10 ppmin comparison to bulk ZnO (T2) and organic-Zn (T3) supplemented fish. But ZnO-NP supplementation beyond 20 ppm did not reveal any added advantage as expression of Cu-Zn SOD gene started todownregulate among the fishes that received the diet fortified with 40 ppm ZnO-NPs. The Cu-Zn SOD gene expression revealed a steady increase upto 20 ppm and further supplementation need to be discouraged.

**Fig 7.**
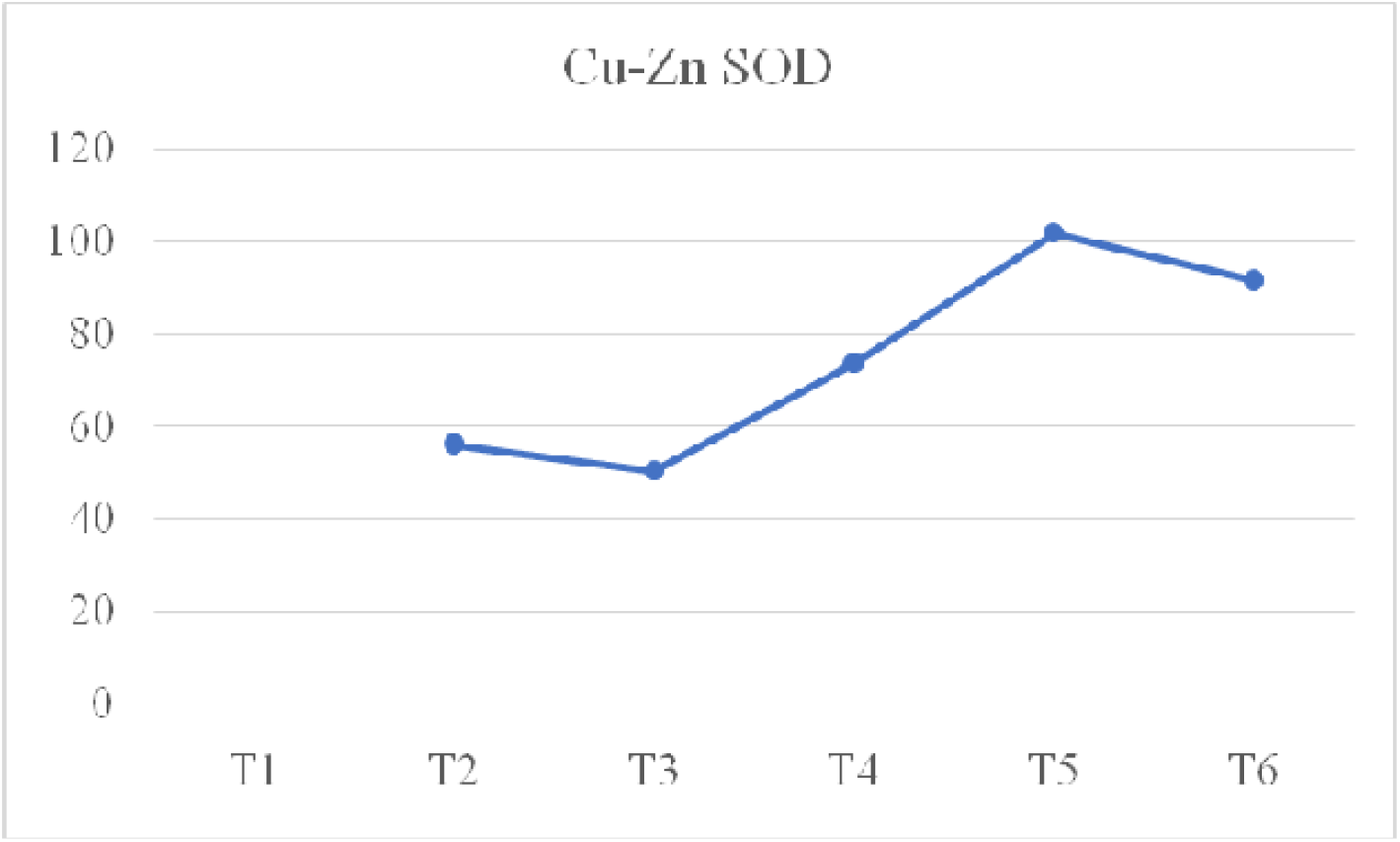
Differential mRNA Expression profiling of Cu-Zn SOD gene of *Chanoschanos* T1 without any added Zinc; T2: Inorganic Zinc 20ppm; T3: Organic zinc 20ppm; T4: Nano zinc 10ppm; T5: Nano zinc 20ppm; T6: Nano zinc 40 ppm.

## Discussion

Zinc is a very important micronutrient needed in required amount at tissue level to impart proper antioxidant status and optimum immunity to fight against oxidative stress. We have synthesized and characterized a unique variety of ZnO-NP to be used for dietary fortification. The bioavailability of as synthesized nano particle is high and having a good anti-microbial potential (23). For dietary fortification the synthesized ZnO-NP was used up to 40 ppm based on the recommendation of our earlier studies (23, 37).

To overcome the issues of bioavailability and to impart higher antioxidant status attempts are initiated **(23, 38, 39**) to device different varieties of nano-zinc to be used for dietary fortification. In a recent study fish diet was fortified with synthesized zinc nano structures (206 nm) and assessed for its positive impact in face of multiple stressors (40). In another attempt different varieties of ZnO-NP (size ranged between 105.7 to 122.4 nm) were devised and evaluated for their efficacy in rohu fish (41). One special variety of dietary ZnO-NP was synthesized having the particle size of 50-60 nm and fortified in the diet of grass carp with an aim for imparting higher body defence (42). In the present set of investigations attempts have been taken to synthesize nano-zinc followed by its microanalytical characterization employing FESEM and PXRD. Once the nanostructure hasbeen confirmed it has been subjected for purity assessment through the Energy Dispersive Spectroscopy. The synthesized ZnO-NPs found to contain only Zinc (71.64%) and Oxygen (28.36%).

In the second phase of this experiment, the antioxidant potential of ZnO-NP was assessed by exploring the Cu-Zn SOD as marker gene. The Cu-Zn SOD is considered to be one of the most important factors in antioxidant activity (44).A number of studies on fish there is limitation on in enzymatic measurement of SOD, which are total or pooled SOD activity without differentiate among the three SOD isoforms, and to transcriptional modulation, which represent one or two distinct fish SODs. These observations do not give a deeper viewof isoform-specificroles of fish SODs. A comparison between the different isoforms of SOD of fish would be important to investigate the different and coordinated mechanisms by which the distinct isoforms protect against oxidative stress (18). That is why we choose Cu-Zn SOD for our present experiment.The antioxidant enzymes like super oxide dismutase and catalase functions to remove the harmful reactive oxygen species (ROS) from cell by activating dissociation of two superoxide radicles to hydrogen per oxide and oxygen.In aquaculture oxidative stress is one of the major threat which often impairs the body defence and hence in turn leads to huge mortality (45).So, there is anurgent need of search for a natural ingredient/ additive to provide a considerable degree of protection by combatting the sudden surge of ROS production.

In the present study nano zinc found to upregulate the expression of Cu-Zn SOD gene. Zinc is the fundamental part of Cu-Zn SOD and a markedly increase in SOD and catalase activity was documented in fish when fed ZnO-NP supplemented diet (46). The fish fed withZnO-NP fortified diet not only exhibited an increased serum antioxidant capacity but also influenced the upregulation of the expression of hepatic SOD, CAT and GPx activity (47).

The KEGG analysis showed that Cu-Zn SOD is active against oxidative stress through longevity regulating pathway. In face of oxidative stress, the Cu-Zn SOD gets activated by the activation of P13K, Akt and dFOXO.

Inorganic zinc, organic zinc and nano-zinc differs in their particle size. The smaller size of nano-zinc provides higher surface area and more efficacy than organic zinc. Due to the bulk size of the other zinc sources (inorganic and organic zinc) remained less available and hence become less interactive with the Cu-Zn SOD protein. The limited bioavailability provides them steric hinderance to fit exactly on the four different zinc binding sites of the protein.

That is why inorganic zinc and organic zinc failed to upregulate the expression of Cu-Zn SOD gene. On the other hand, nano-zinc being dwarfand less interactive with other nutrients, seems to be more bioavailable andeasily get attachedto the protein and binds with the different zinc binding sites of the protein. When nano-zinc binds with the Cu-Zn SOD protein, it activates the gene in face of oxidative stress and therebyupregulates its expression. With the increasing level of dietary fortification there is more available zinc at cellular level which ensures more zinc to crosstalk with the protein and influencing for its better expression. But it remained true till the dietary fortification level up to 20 ppmZnO-NP, beyond which no more upregulation of Cu-Zn SOD was observed. The fish fed diet with 40ppm ZnO-NP fortification did not reveal any added advantage. Hence from the resent study it appeared that dietary supplementation of 20 ppm ZnO-NPs is best for maximizing the Cu-Zn SOD expressionand in turn providing the highest antioxidant activity of the cell.

## Conclusion

In the present study, the synthesis of nano zinc oxide particles through colloidal chemistry method was successful, and their nano size was confirmed by the analysis of micro-analytical structures. Dietary fortification of fish with nano zinc revealed the highest Cu-Zn SOD expression. The higher availability of nano-zinc at cellular level allowed it to saturate the zinc binding sites of Cu-Zn SOD protein and in turn can modulate its expression profile. The upregulation of the Cu-Zn SOD appears to be dose dependent and the highest influence was observed when the feed was fortified with at 20 ppm ZnO-NP beyond which no more added advantage was recorded. This is the first report to document the influence of dietary nano-zinc in the expression profiling of Cu-Zn SOD of *Chanos chanos*. In this study we explored the mechanism of action of *nano zinc* improving the antioxidant status with *Cu-Zn SOD gene as molecular marker* and *Chanos chanos* as model organism.

