## Supplementary figures and images for "Supplemental Dietary Nano Zinc Can Impart Higher Antioxidant Status by Modulating Cu-Zn SOD Expression"

### Suppl1

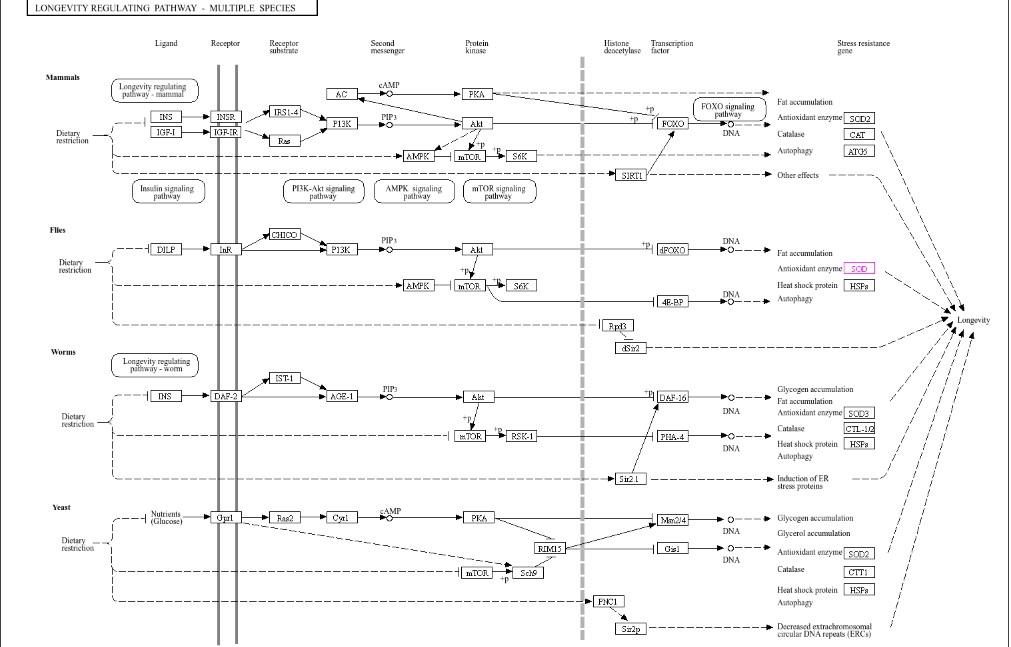
